# Integrating Unsupervised Clustering with Transformer-Based Models for De Novo Cell Population Identification

**DOI:** 10.64898/2026.11.12.766182

**Authors:** Yanting Guo, Guangyou Duan, Guofeng Xu, Chang Liu, Jia Chang, Yangguang Han, Xin Li, Shan Gao

**Affiliations:** College of Life Sciences, Nankai University, Tianjin, 300071, P.R.China; School of Mathematics and Statistics, Yunnan University, Kunming, Yunnan 650504 China; School of Life Sciences, Qilu Normal University, Jinan, Shandong 250200, P.R.China; School of Medicine, Nankai University, Tianjin 300071, P.R.China

**Keywords:** Artificial Intelligence, scRNA-seq, 10x Genomics, Smart-seq2, biomarker discovery

## Abstract

Single-cell transcriptome sequencing has been widely applied across the life sciences. It enables accurate and robust identification of cell types in heterogeneous biological tissues through unsupervised clustering, shallow learning, and deep learning. While deep learning does not outperform shallow learning in classification performance, it streamlines cell type identification by eliminating cumbersome preprocessing procedures. However, the availability of training datasets with comprehensive cell type annotations is always limited, so new datasets inevitably contain cell populations that are not pre-defined in the training data. To address this limitation, we integrated unsupervised clustering with transformer-based models to identify cell populations unrecognized by classical methods. Specifically, we developed the Nankai GeneRank Classifier by fine-tuning the pre-trained Geneformer-V1 model, and then clustered cells based on its softmax-normalized class probability matrix (SCPM). Applied to a public colorectal cancer dataset, the new method, named SCPM-based clustering, resolved cell populations that cannot be unambiguously identified by classical methods alone and led to the discovery of novel candidate markers of colorectal cancer stem cells, demonstrating its utility.

## Introduction

Single-cell transcriptome sequencing [1], a transformative technology, has been widely applied across the life sciences and enables accurate and robust identification of cell types in heterogeneous biological tissues. A broad range of mainstream methods have been developed for this purpose, spanning unsupervised clustering, shallow learning, and deep learning. Representative tools in these three categories include the Seurat package [2] for unsupervised clustering, CellTypist [3] for shallow learning, and Geneformer [4] for deep learning. As the most widely used single-cell analysis tool, the Seurat package integrates principal component analysis (PCA), K-nearest neighbor (KNN) graph construction, shared nearest neighbor (SNN) computation, Leiden or Louvain clustering and Uniform Manifold Approximation and Projection (UMAP [5]) visualization into a unified pipeline to facilitate cell type identification. CellTypist is built on logistic regression optimized via stochastic gradient descent and performs one-vs-rest binary classification of cell types, serving as a representative shallow-learning framework. In contrast, Geneformer adopts a BERT-style [6] transformer [7] architecture pre-trained on millions of single-cell transcriptomes and requires supervised fine-tuning for downstream cell classification tasks.

In our previous studies, we observed that almost all tested shallow- and deep-learning methods achieved classification accuracies above 90%, exhibiting the high intrinsic class separability of single-cell transcriptomic datasets (*i*.*e*., scRNA-seq dataset). In parallel, unsupervised clustering results showed high concordance with those of classification methods. However, both unsupervised clustering and shallow-learning methods rely on cumbersome preprocessing steps, *e*.*g*., data normalization [8] and PCA dimensionality reduction. Moreover, cell type identification using unsupervised clustering is highly sensitive to hyperparameter perturbations—including clustering resolution and nearest-neighbor graph construction settings—and depends heavily on the manual selection of marker genes, which is inherently subjective. In contrast to these two paradigms, deep-learning methods, particularly transformer-based models, offer a wide range of advantages beyond eliminating cumbersome preprocessing. For instance, they can readily process heterogeneous data (e.g., 10x Genomics and Smart-seq2) without requiring data integration. Nevertheless, all of these methods inherently rely on prior knowledge of known cell types or their markers, and the development of new methods or strategies capable of discovering novel cell types or subtypes remains the most promising future direction for the field.

In the present study, we developed the Nankai GeneRank Classifier, a BERT-style transformer-based model for cell type identification using scRNA-seq data. Built upon the pre-trained Geneformer framework, the classifier was fine-tuned on a dataset comprising 29 pre-defined cell types with a total of 54,362 cells. We then integrated unsupervised clustering with the classifier’s softmax-normalized class probability matrix (SCPM) to identify cell populations unrecognized by classical methods, yielding the SCPM-based clustering. Applied to a public colorectal cancer dataset, this method resolved previously unrecognized cell populations and identified novel candidate markers, extending the findings reported in the original study [9].

## Results

### Performance comparison of shallow and deep learning-based cell classification models

As a foundational observation preceding the development of our new method, almost all tested shallow- and deep-learning methods achieved classification accuracies above 90%, underscoring the high intrinsic class separability of scRNA-seq datasets (**Introduction**). To illustrate this point, we employed a high-quality scRNA-seq dataset (GEO: GSE188515) generated from our previous study [10], in which 16 distinct cell types were identified from 10,784 cells through Seurat clustering (**Materials and Methods**). Using this dataset, we benchmarked Geneformer-based classification models against nine widely used shallow-learning methods, as well as against the well-established tool CellTypist (**Introduction**), which is likewise based on shallow learning. The nine methods [11] used to construct classification models included Support Vector Machine (SVM), Random Forest (RF), Generalized Boosted Model (GBM), Recursive Partitioning and Regression Tree (RPART), Artificial Neural Network (ANN), Partial Least Squares (PLS), K-Nearest Neighbor (KNN), Naive Bayes (NB), and AdaBoost.

To guarantee unbiased benchmarking, we applied a unified data preprocessing workflow across all shallow learning methods, whereas Geneformer-based classification models require only a single preprocessing step: extraction of the top 2,048 genes ranked by their expression levels. The performance of all methods was evaluated using accuracy and Matthews Correlation Coefficient (MCC) under standard 10-fold cross-validation (**Materials and Methods**). All shallow-learning models performed on par with the Geneformer-based classification models (**Table 1**), although the former require cumbersome preprocessing steps (**Introduction**). This may be attributed to the facts that classification performance directly depends on the statistical relationship between features and pre-defined classes, and the consistently high performance across heterogeneous methods and data types reflects a certain degree of causal association between cell types and gene expression ranks. Notably, when the Geneformer-based classification model was randomly initialized and fine-tuned on the 10,784 cells without pre-training, it achieved an accuracy of 0.9334 (MCC 0.9239), slightly lower than the 0.9609 (MCC 0.9553) obtained by fine-tuning the model initialized with weights pre-trained on ∼30 million cells. Thus, pre-training was not required for achieving high accuracy.

**Table 1.** Performances on all models.

| Model | 10 cross-validation |  | Optimized hyperparameters |
|---|---|---|---|
|  | Accuracy | MCC |  |
| CellTypist | 0.9587 | 0.9528 | C=1 |
| SVM-Linear | 0.9645 | 0.9594 | C=10 |
| RF | 0.9610 | 0.9553 | mtry=39, ntree=300 |
| GBM | 0.9621 | 0.9566 | interaction.depth=10<br>n.trees=250, shrinkage=0.1 |
| RPART | 0.8173 | 0.7899 | Maxdepth=10 |
| ANN | 0.9688 | 0.9643 | Size=50, decay=0.1 |
| PLS | 0.9330 | 0.9235 | Ncomp=30 |
| KNN | 0.9670 | 0.9622 | K=10 |
| NB | 0.9414 | 0.9330 | fL=0.01, usekernel=TRUE |
| AdaBoost | 0.9538 | 0.9472 | maxdepth=5, iter=250, nu=0.3 |
| Geneformer pre | 0.9609 | 0.9553 | learning rate=2e-4, epochs=10 |
| Geneformer * | 0.9334 | 0.9239 | learning rate=2e-4, epochs=10 |

Model performance was quantified using accuracy and Matthews Correlation Coefficient (MCC). Based on the GSE188515 dataset, the performance of all models was evaluated with their optimized hyperparameters setting, which were determined through an exhaustive grid search with 10-fold cross-validation. The difference between Geneformer pre and Geneformer \* mainly stems from their initial weight matrices: the initial weight matrix of Geneformer pre is derived from a pre-trained model, while that of Geneformer \* is randomly generated.

### Nankai GeneRank Classifier and its application

Built upon the pre-trained Geneformer-V1 architecture, we first performed supervised fine-tuning on the fully annotated GSE188515 dataset to develop the Nankai GeneRank Classifier v1. Starting from the v1 checkpoint, we further conducted incremental fine-tuning by integrating three additional public scRNA-seq datasets: 1,160 cells of 3 cell types from GSE232996 [12], 3,100 cells of 1 cell type from GSE180286 [12], and 39,318 cells of 9 cell types from GSE162610 [13]. All four datasets were generated using the 10x Genomics scRNA-seq technologies. Specifically, GSE188515 and GSE180286 are derived from human samples, while GSE232996 and GSE162610 correspond to mouse transcriptomic data. The feasibility of jointly training on combined human and mouse data from different tissues rests on our previous findings that differences across species, tissue origins, and library-construction and sequencing technologies exert a negligible impact on the cell-type identification accuracy of our model, a conclusion further supported by the present case. The multi-dataset training yielded the v2 classifier, which encompasses a total of 54,362 cells across 29 distinct cell types (**Supplementary File 1**). Notably, three of the 29 cell types—endothelial cell type 2 (EC2), ependymal cell, and breast cancer cell—originated from pediatric cavernous hemangioma [10], mouse spinal cord [13], and human breast cancer samples [12], respectively. EC2 denotes endothelial cells exhibiting a certain degree of abnormality [10], whereas ependymal cell is the closest counterpart to epithelial cell (EpC), which was not included in the training set. Given their high similarity in gene expression ranks to their respective closely related cell types, EC2, ependymal, and breast cancer cells can be used as reference to detect ECs, EpCs, and cancer cells, respectively.

The Nankai GeneRank Classifier v2 was tested on a public Smart-seq2 dataset cc-scRNA783 (**Materials and Methods**), which comprises 783 single cells isolated from tumor and adjacent tissues of colorectal cancer patients [9]. Each of the 783 cells was then assigned to one of 29 pre-defined cell types using the classifier. Testing a model trained on 10x Genomics data against a Smart-seq2 dataset constitutes a cross-platform validation, with important implications for single-cell transcriptome integration and downstream functional interpretation. Smart-seq2 and 10x Genomics constitute two representative scRNA-seq technologies [1]. Smart-seq2 achieves superior gene detection sensitivity but captures fewer cells overall due to its elevated per-cell cost. Conversely, 10x Genomics facilitates profiling of substantially larger cell numbers, albeit with a considerably lower median number of detected genes per cell. Consequently, minor misclassifications introduced by clustering algorithms are largely negligible in 10x Genomics datasets encompassing tens of thousands of cells, as their impact is diluted across the expansive cell population. In Smart-seq2 datasets, however, where cell numbers are inherently constrained, such errors become drastically amplified: even a handful of misassigned cells contaminating a small cluster can severely compromise marker gene detection, culminating in ambiguous or erroneous cell type identification. In the subsequent analysis, we compare Seurat clustering results with those predicted by Nankai GeneRank Classifier v2, which serves as a unified representative for all included shallow- and deep-learning classification methods (**Table 1**).

Using Seurat, the 783 cells were clustered into nine clusters (**Materials and Methods**), designated as o0–o8, where o denotes original (**Figure 1A**). To identify these clusters, we mapped each cell to the 29 pre-defined cell types according to its binary prediction (0 or 1) from the Nankai GeneRank Classifier. The similarity between each cluster (o0–o8) and each cell type was subsequently quantified as the proportion of cells within that cluster assigned to the corresponding cell type (**Figure 1B**). For instance, the similarities between o1 and EC2, CD4+ TC, CD8+ TC, and BC were 0.08, 0.50, 0.24, and 0.09, respectively. Based on these cluster–cell-type similarity profiles, o1, o2, o8, o4, and o7 were assigned to T cell (TC), B cell (BC), mast cell, M1-like macrophage (m1Maph), and M2-like macrophage (m2Maph), respectively, consistent with the findings reported in the original study [9]. Among the nine clusters, o5 was uniquely characterized by its highest similarity to breast cancer cell, suggesting its correspondence to the previously reported cancer stem cells (CSCs) [9]. Each of the remaining three clusters (o0, o3, and o6) exhibited high similarity to multiple cell types (**Figure 1B**), indicative of potentially unrecognized cell populations. We therefore concluded that: (1) Seurat clustering and the classifier achieved substantial yet incomplete concordance; and (2) the three unrecognized clusters could not be unambiguously identified by classical classification methods alone.

**Figure 1.**
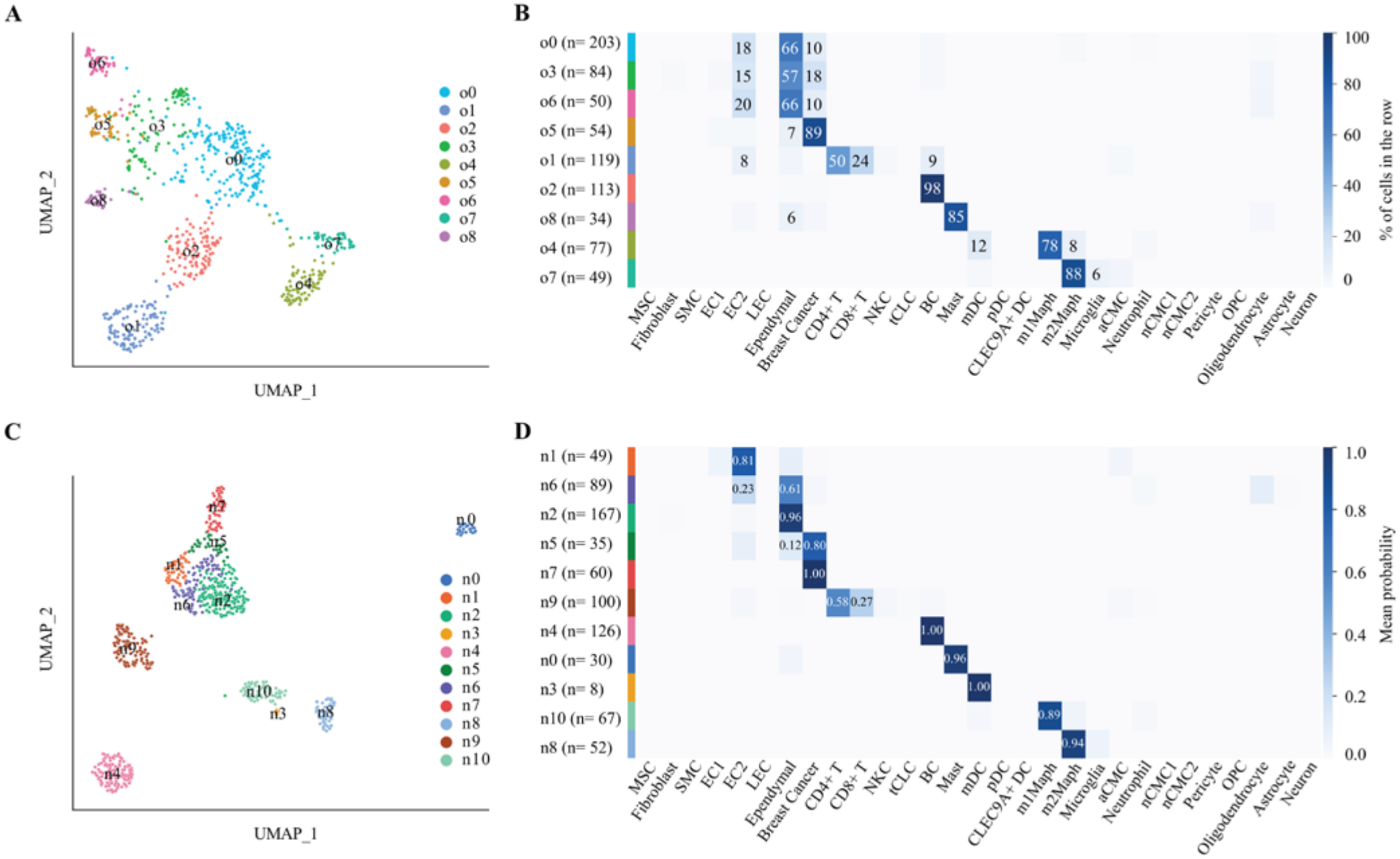
Understanding the cluster-cell-type similarities. The cc-scRNA783 dataset comprises 783 single cells isolated from tumor and adjacent tissues of colorectal cancer patients (Materials and Methods). A. 783 cells were clustered into nine clusters, designated o0-o8, using Seurat clustering. B. Each cell (y-axis) was mapped to 29 pre-defined cell types (x-axis) according to its binary prediction (0 or 1) from the Nankai GeneRank Classifier and the similarity between each cluster and each cell type was subsequently quantified as the proportion of cells within that cluster assigned to the corresponding cell type. Values and colors are both shown when the proportion exceeds 0.05; only the color is displayed otherwise. C. 783 cells were clustered into 11 clusters, designated n0-n10, using softmax-normalized class probability matrix (SCPM)-based clustering. D. Each cell (y-axis) was mapped to 29 pre-defined cell types (x-axis) according to its corresponding row in the SCPM and the similarity profiles between each cluster and 29 cell types were subsequently quantified by averaging the SCPM rows of all cells (i.e., mean probabilities) within the cluster. Values and colors are both shown when the mean probability exceeds 0.05; only the color is displayed otherwise.

### Development of new method to identify unrecognized cell populations

Because pairwise similarity between each query cell and the pre-defined cell type is inherently encoded in the softmax-normalized class probability matrix (SCPM), our new method performs downstream clustering directly on this matrix, thereby enabling cell-type identification at the cluster level rather than relying on single-cell-level binary predictions from the Nankai GeneRank Classifier. In contrast to Seurat clustering, which operates on PCA results, our method relies on the SCPM and is hereafter referred to as SCPM-based clustering. The two-step workflow of SCPM-based clustering comprises SCPM generation via the Nankai GeneRank Classifier, followed by a clustering step that integrates centered log-ratio matrix transformation, KNN graph construction, and Leiden community detection. Here, we use the cc-scRNA783 dataset as a representative example to illustrate this workflow. Consequently, SCPM-based clustering yielded 11 distinct clusters (**Figure 1C**), designated n0–n10, where n denotes new.

To identify these new clusters, we mapped each cell to the 29 pre-defined cell types according to its corresponding row in the SCPM. The similarity profile between each cluster (n0–n10) and the 29 pre-defined cell types was subsequently quantified by averaging the SCPM rows of all cells (*i*.*e*., mean probabilities) within the cluster (**Figure 1D**). Based on the cluster–cell-type similarity profiles, six clusters were unambiguously identified as the six cell types (CSC, TC, BC, mast cell, m1Maph, and m2Maph), consistent with the cell types reported in the original study [9]. Comparative analysis of the original clusters (**Figure 1B**) and new clusters (**Figure 1D**) against the 29 pre-defined cell types revealed that o0, o3 and o6 correspond to n1, n6, n2 and n5, representing the unrecognized cell populations. These unrecognized cell populations were similar to three pre-defined cell types—EC2, ependymal cell, and breast cancer cell—which correspond to the EC, EpC, and CSC identified in the original study [9]. However, the similarity profiles of the original clusters o0, o3 and o6 across the three pre-defined cell types failed to exhibit significant differences, thereby precluding unambiguous identification. If assignment were based on the highest similarities (66% for o0, 57% for o3, and 66% for o6), they would all be classified as EpC, given that the ependymal cell type represents EpC. In contrast, the four new clusters exhibited clearly distinct similarity profiles, underscoring the higher resolution achieved by SCPM-based clustering.

Under the classical analytical framework, which employs the highest similarities for cell-type assignment, n1, n6, n2, n5, and n7 would necessarily be collapsed into three clusters (n1, n6&n2, and n5&n7), corresponding to three pre-defined cell types (EC, EpC, and cancer). In contrast, SCPM-based clustering revealed that these five clusters actually corresponded to three pre-defined cell types (EC, EpC, and cancer) as well as two intermediate cell types, as evidenced by their distinct similarity profiles. Accordingly, we designated them as ccEC, ccEpC, malignant ccEpC, cancer-like ccEpC, and CSC, respectively, where “cc” denotes cells derived from colorectal cancer tissue. These five cell types recapitulate a graded spectrum of similarity to the pre-defined cell types, spanning from EC through EpC to the cancerous state (**Figure 1D**). Therefore, our new method successfully resolved two previously unrecognized cell populations—malignant ccEpC and cancer-like ccEpC—that were missed in the original study [9].

### Advantages of SCPM-based clustering

SCPM-based clustering has two key advantages: it enhances cell type identification resolution by using clustering instead of the classical maximum-mean-probability assignment criterion, and it ensures accuracy (**Table 1**) through deep learning optimization by Nankai GeneRank Classifier instead of Seurat clustering. Despite broad global concordance, notable compositional differences were observed between the corresponding original/new cluster pairs o5/n7, o1/n9, o2/n4, o8/n0, o4/n10, and o7/n8 (**Figure 2**). In particular, SCPM-based clustering identified an additional cluster n3 as mDC that was missed by Seurat clustering. This mDC cluster consists of only eight cells. According to the Seurat clustering results, seven of these eight cells were assigned to m1Maph and the remaining one was assigned to m2Maph. Moreover, in the UMAP plot (**Figure 2**), cells with discordant assignments between the two methods were not exclusively located at cluster boundaries, indicating that these discrepancies cannot be resolved simply by hyperparameter adjustment. Further compelling evidence for this conclusion is provided by the complete breakdown of mapping correspondence between the three original clusters (o0, o3, and o6) and the four new clusters (n1, n6, n2, and n5). Given that the CSC cluster was rigorously validated in the original study [9], we performed Seurat clustering for comparative analysis between the n7 cluster and its corresponding counterpart, via exhaustive gridding of hyperparameters including the number of highly variable genes, principal components, nearest neighbors, clustering algorithm, and clustering resolution (**Supplementary File 1**). Across all resulting clustering outputs, the cluster exhibiting the highest similarity to n7 contained 53 cells, achieving a Jaccard index of merely 0.662 with only 45 overlapping cells. When hyperparameters were improperly specified, CSCs failed to form a distinct cluster and instead aggregated with other cells into larger clusters, the largest of which encompassed up to 432 cells. Collectively, these findings demonstrate that the accuracy of cell-type identification in SCPM-based clustering can not be ensured by classical clustering.

**Figure 2.**
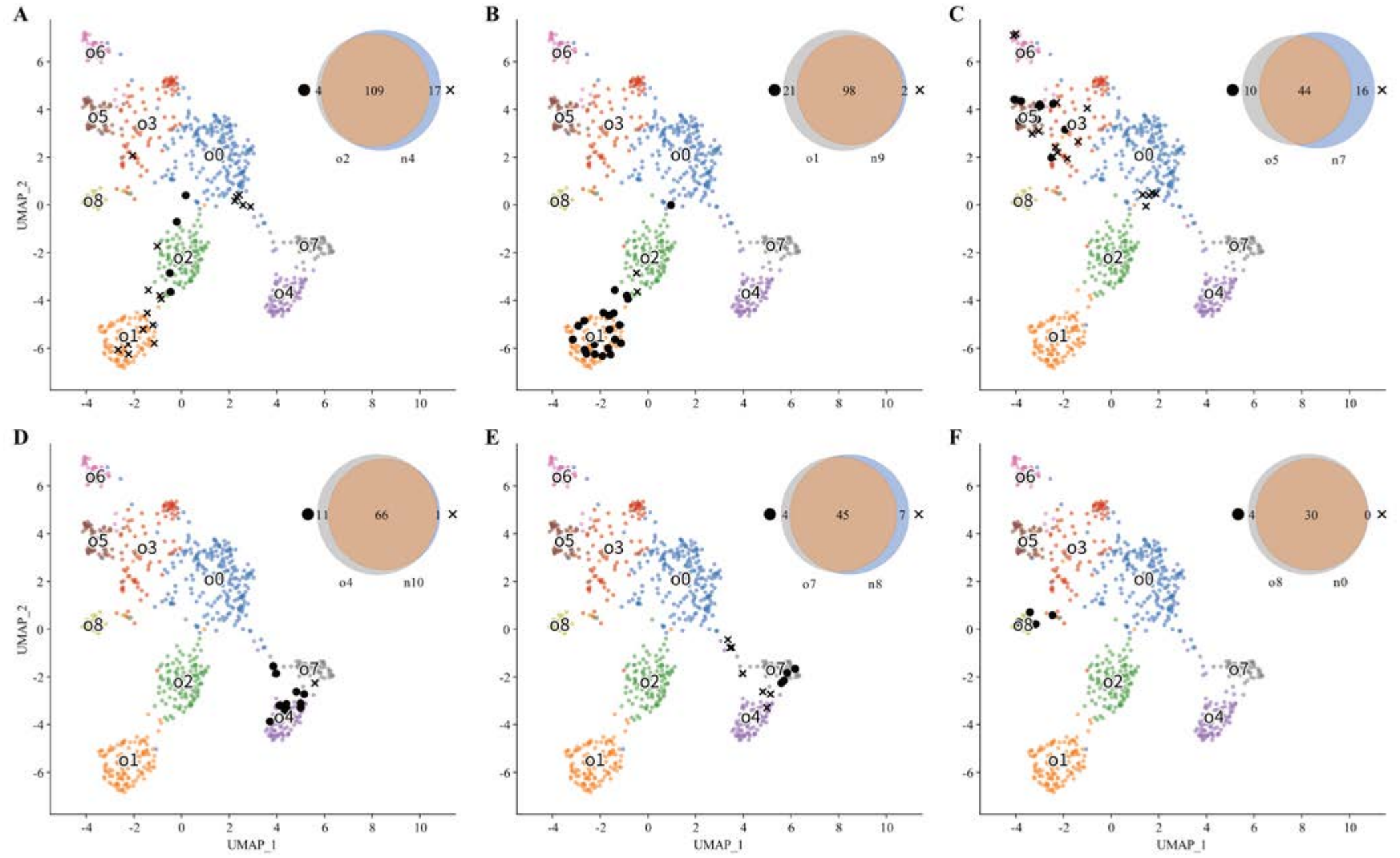
Comparison of the original clusters and new clusters. 783 cells were clustered into nine clusters, designated o0-o8 (Figure 1A), using Seurat clustering and into 11 clusters, designated n0-n10, using softmax-normalized class probability matrix (SCPM)-based clustering (Figure 1C). This UMAP plot is a direct replicate of the clustering layout shown in Figure 1A, and cells with discrepant cluster assignments between the two methods are marked by black solid circles and crosses.

As an additional key advantage, SCPM-based clustering enables the unified processing of both bulk RNA-seq and scRNA-seq datasets for direct cross-modal comparison. Based on the pairwise similarity between each query cell and the pre-defined cell type, SCPM-based clustering derives cluster-cell-type similarities. By treating each bulk sample as an independent “pseudo-cluster”, we further extend this framework to compute tissue-to-cell-type similarity, enabling direct comparison between bulk-derived and single-cell-derived cell-type signatures (**Figure 3**). As a proof of this concept, we applied a bulk RNA-seq dataset experimentally matched to the cc-scRNA783 dataset. This dataset comprises nine pairs of histologically confirmed samples from colorectal cancer patients, with each pair consisting of matched tumor and adjacent tissues. In the UMAP plot (**Figure 3A**), the nine tumor samples clustered closer to the CSC cluster than did the nine adjacent samples. Mapping these samples onto the 29 pre-defined cell types further highlighted differences between the two groups (**Figure 3B**): overall, similarity to EC2 was detected in almost all samples, whereas similarity to EpC or cancer cell types was detected in five tumor samples but only one adjacent sample. Unexpectedly, similarity to oligodendrocytes was also detected in both tumor and adjacent samples, which merits further in-depth investigation.

**Figure 3.**
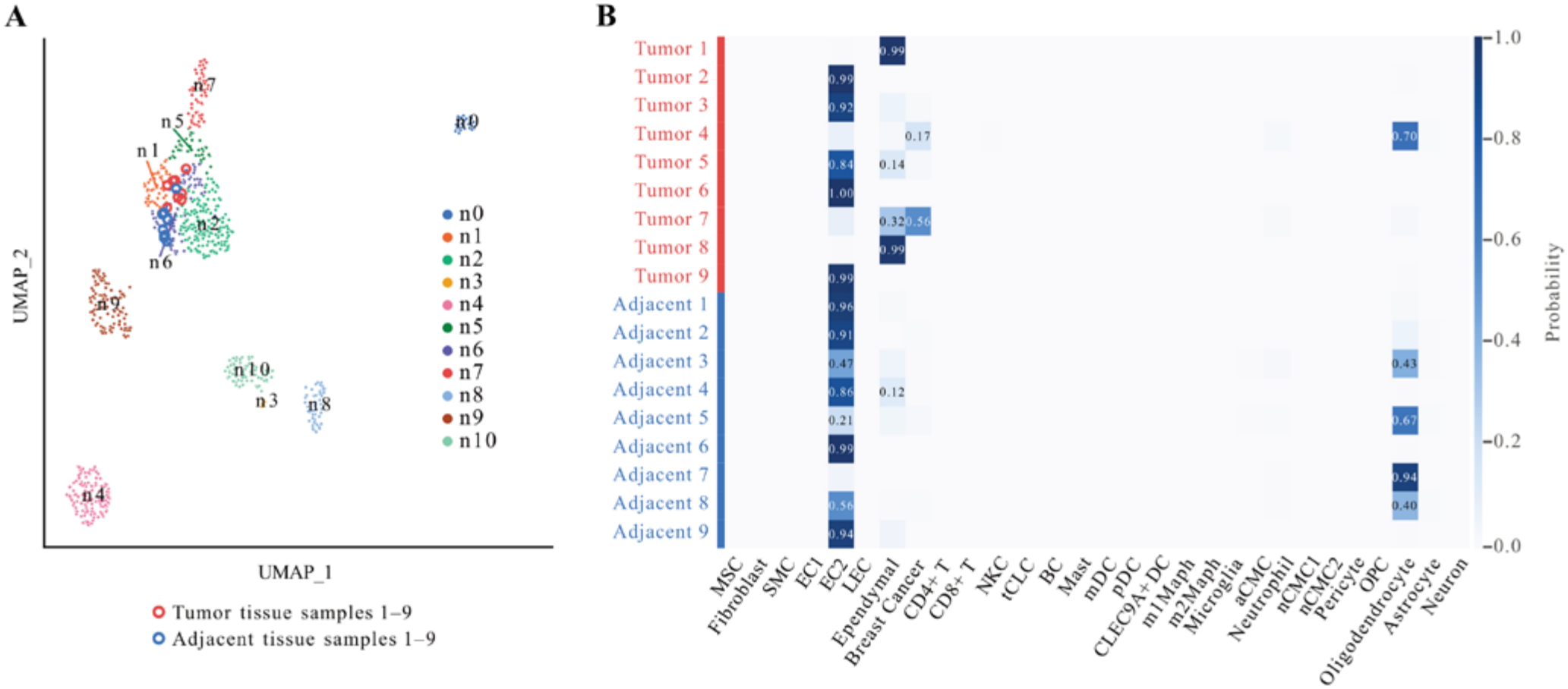
Comparison of single cells and bulk samples. A. Each bulk sample was mapped into this UMAP embedding according to its nearest single-cell neighbors in clr space, and the cluster labels and UMAP coordinates of the single cells were unchanged as shown in Figure 1C. B. Each bulk sample (y-axis) was mapped to 29 pre-defined cell types (x-axis) according to its corresponding row in the SCPM and the similarity profile between bulk sample and the 29 cell types was subsequently quantified by its corresponding row. Values and colors are both shown when the probability exceeds 0.05; only the color is displayed otherwise.

### Discovery of novel potential CSC markers

Based on the cc-scRNA783 dataset, the original study [9] reported that no single previously published marker could effectively distinguish colorectal CSCs from other cell populations. These markers, including PROM1/CD133, CD44, LGR5, SSEA4 (a glycosphingolipid antigen), BMI1, LRIG1, KRT18, SLC12A2, CXADR, and CLDN4 (**Figure 4**), among which CD133, CD44, LGR5 and SSEA4 were used in the original study [9] to enrich CSCs by fluorescence-activated cell sorting (FACS). However, FACS enrichment analysis showed that the cell types with the highest enrichment proportion for LGR5 were m1Maph and m2Maph, those for CD44 were TC, mast cell, and BC, whereas that for PROM1 was EpC (**Figure 4A**). In addition, the present study demonstrated that CSCs, as a special type of cycling cells [12], exhibit high expression of 26 markers that clearly distinguish them from the other 10 cell types (**Figure 4B**). Accordingly, the elevated expression of these 26 markers provides compelling evidence supporting the CSC identity of cluster n7. Further evidence comes from the observation that 46 of the 81 cells (57%) derived from sphere culture were assigned to the n7 cluster.

**Figure 4.**
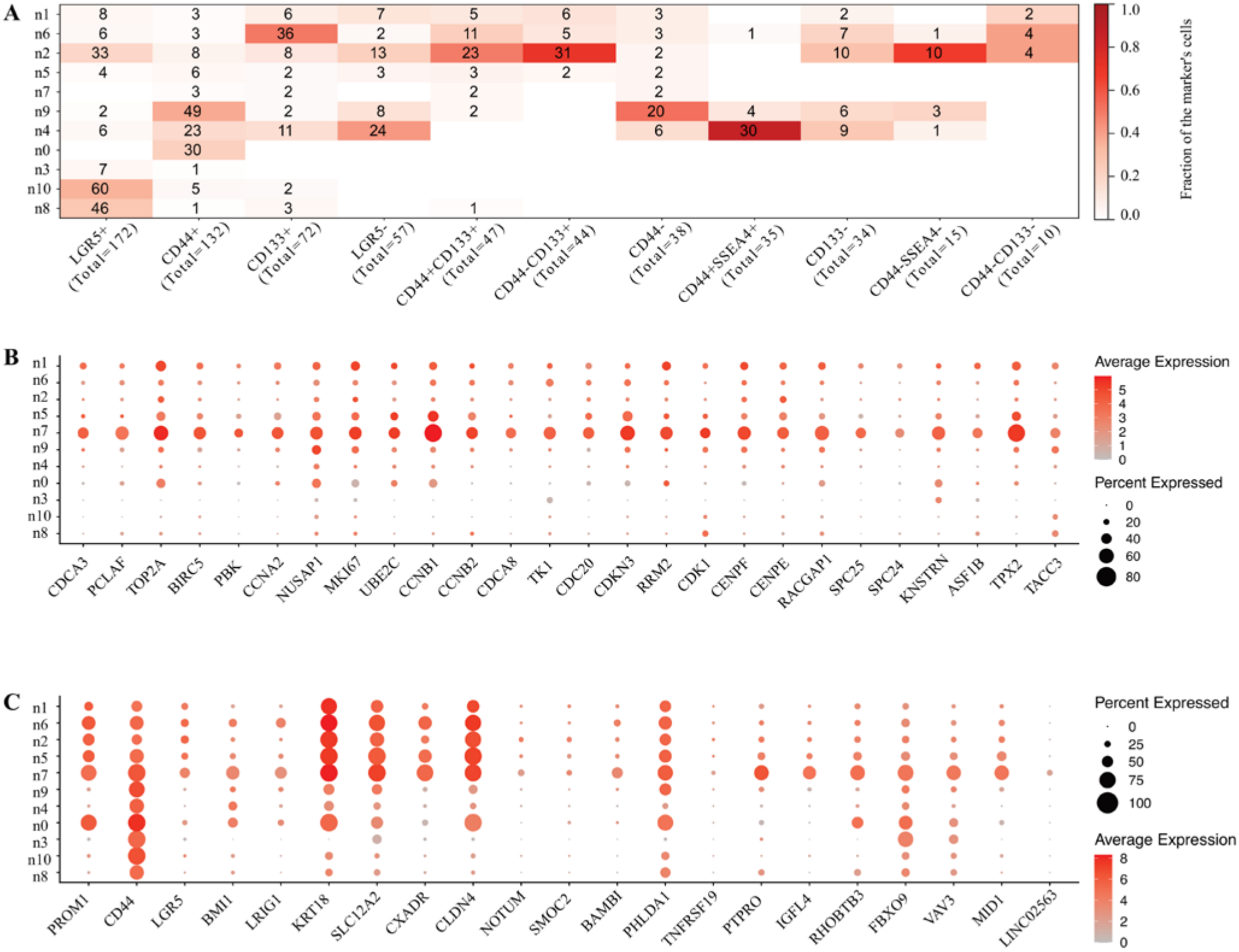
Marker gene expression across cell types. 783 cells were clustered into 11 clusters, designated n0-n10, using softmax-normalized class probability matrix (SCPM)-based clustering. Clusters n0-n10 were identified as mast cell, ccEC, malignant ccEpC, mDC, BC, cancer-like ccEpC, ccEpC, CSC, m2Maph, TC, and m1Maph. A. In the original study [9], PROM1/CD133, CD44, LGR5, SSEA4 (a glycosphingolipid antigen) were used as markers for CSC isolation in FACS experiments. B. A set of 26 marker genes can be used to identify CSCs, as they represent a special type of cycling cells [12]. C. Early and recently reported CSC marker genes are compared with the marker genes identified in the present study.

To discover potential CSC markers based on SCPM-based clustering, we next sought to identify protein-coding genes and long non-coding RNAs (lncRNAs) more specifically expressed in CSCs than in the other 10 cell types. The most common approach selects candidates from significantly differentially expressed (DE) genes between CSC and non-CSC groups (**Supplementary File 2**); using this approach, the previous study [9] recently reported five CSC marker genes (*NOTUM, SMOC2, BAMBI, PHLDA1*, and *TNFRSF19*). In contrast, based on our re-identified five cell types, we found 370 genes and lncRNAs (**Supplementary File 1**) whose expression progressively increased from ccEC and ccEpC, through malignant ccEpC and cancer-like ccEpC, to CSCs. Among these, 164 showed a gradual increase, whereas the remaining 206 exhibited a sharp rise at the final step, *i*.*e*., a substantially higher fold change between cancer-like ccEpC and CSCs (**Figure 5**). Of the 164 genes and lncRNAs, only 46 overlapped with the DE genes, and all except DST were expressed at very low overall levels. By comparison, 150 of the 206 genes and lncRNAs overlapped with the DE genes. Notably, none of the five recently reported marker genes [9] overlapped with either the 164 or the 206 gene set.

**Figure 5.**
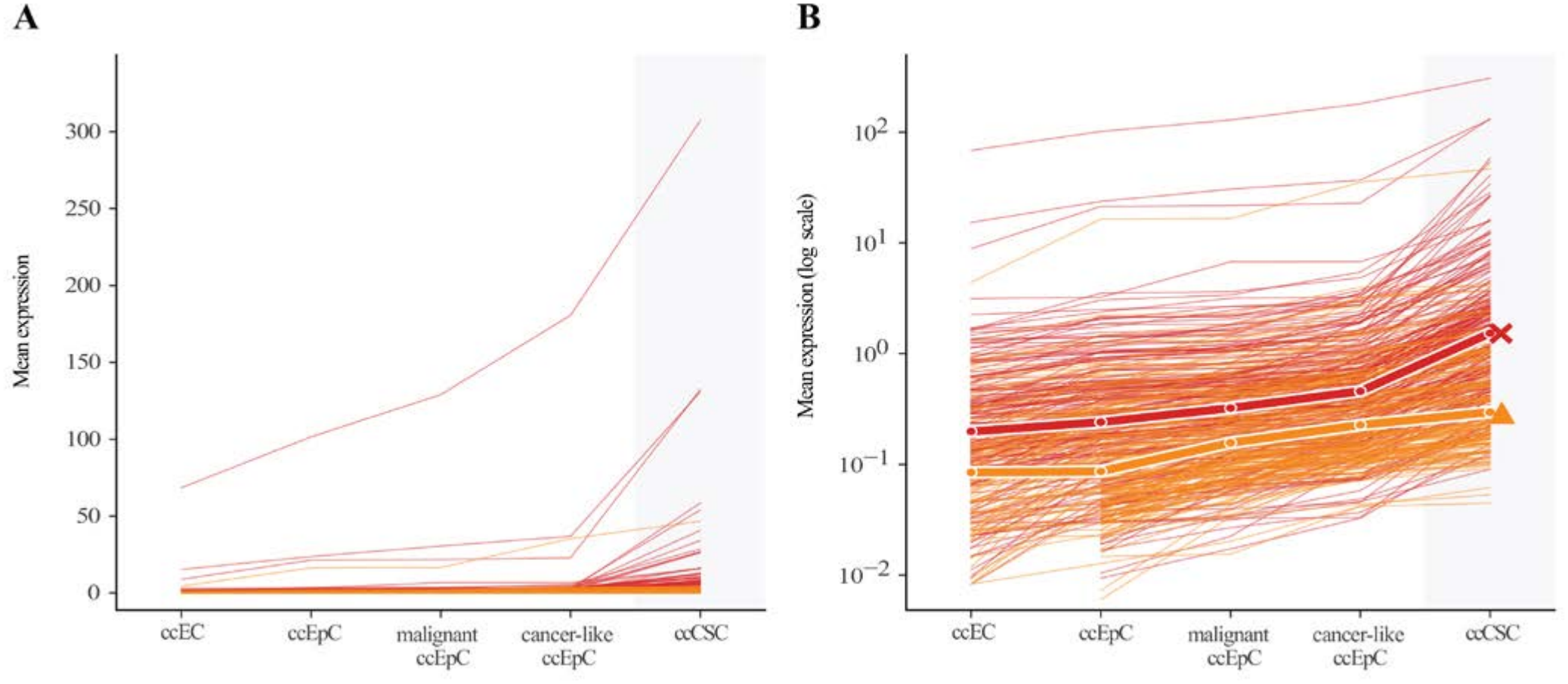
Two patterns of gene expression up-regulation across five cell types. 783 cells were clustered into 11 clusters, designated n0-n10, using softmax-normalized class probability matrix (SCPM)-based clustering. Clusters n1, n6, n2, n5, and n7 were identified as ccEC, ccEpC, malignant ccEpC, cancer-like ccEpC, and CSC (x-axis), respectively, where “cc” denotes cells derived from colorectal cancer tissue. A. 164 (in orange color) exhibited a gradual increase in mean expression (y-axis) across the five cell types (x-axis), whereas the remaining 206 (in red color) displayed a sharp rise at the final step, i.e., a substantially higher fold change between cancer-like ccEpC and CSCs. B. The same data as in A is shown on a base-10 logarithmic y-axis and the medians of the 164 and 206 molecules are indicated by triangles and crosses, respectively

Among the 150 overlapping genes and lncRNAs, the lncRNAs showed substantially lower expression levels than the protein-coding genes. We therefore ranked all 150 molecules by their fold changes between cancer-like ccEpCs and CSCs. The top six were protein-coding genes (*PTPRO, IGFL4, RHOBTB3, FBXO9, VAV3*, and *MID1*), with fold changes of 26.3, 17.3, 15.5, 15.0, 13.0, and 12.4, respectively, and the seventh was the lncRNA LINC02563, with a fold change of 11.3. Notably, *PTPRO* was first reported to repress colorectal cancer tumorigenesis [14], and subsequent work proposed that PTPRO-high stem-like cells interact with CAFs and endothelial cells through the DLL4–NOTCH signaling axis [15]. In addition, circRHOBTB3, a circular RNA derived from the *RHOBTB3* locus, is secreted via exosomes to sustain colorectal cancer cell fitness [16]. The marker–cluster dot plot showed that *PTPRO, IGFL4, MID1* [17], and LINC02563 [18] exhibited greater CSC specificity than *RHOBTB3, FBXO9, VAV3*, and the five recently reported *NOTUM, SMOC2, BAMBI, PHLDA1*, and *TNFRSF19* (**Figure 4C**). However, LINC02563 was detected in only 28% of CSCs, attributable to its substantially lower expression—on average only 1.37% of that of *PTPRO*.

## Materials and methods

### Data acquisition, dataset construction and sample encoding

Four public 10x Genomics scRNA-seq datasets, including GSE188515, GSE162610, GSE232996 and GSE180286, were downloaded from the NCBI GEO database. The colorectal cancer dataset initially comprised 831 single cells isolated from tumor and adjacent tissues of colorectal cancer patients generated by Smart-seq2 in the original study [9]. Its raw FASTQ files are publicly accessible in the NCBI SRA database under the accession number SRP113436. While the original study [9] utilized only a final high-quality dataset of 693 cells, the present study refined the data preprocessing pipeline, yielding an improved dataset of 783 cells, designated cc-scRNA783 (**Supplementary File 1**). The 29 pre-defined cell types included Fibroblast, mesenchymal stem cell (MSC), smooth muscle cell (SMC), EC1, EC2, lymphatic endothelial cell (LEC), CD4 positive T cell (CD4+ TC), CD8 positive T cell (CD8+ TC), natural killer cell (NKC), BC, mast cell, monocyte derived dendritic cell (mDC), plasmacytoid dendritic cell (pDC), CLEC9A positive dendritic cell (CLEC9A+ DC), m1Maph, m2Maph, pericyte, oligodendrocyte precursor cell (OPC), oligodendrocyte, astrocyte, ependymal cell, neuron, cycling myeloid cell type a (aCMC), microglia, neutrophil, cycling lymphoid cell type t (tCLC), cycling myeloid cell type n subtype 1 (nCMC1), cycling myeloid cell type n subtype 2 (nCMC2), and breast cancer. The detailed information of the 29 pre-defined cell types is provided in **Supplementary File 1**.

### Model training, tuning and data processing

SVM, RF, GBM, RPART, ANN, PLS, KNN, NB, and AdaBoost were briefly described in our previous study [11]. After evaluating SVM methods with two distinct kernel functions, SVM with a linear kernel (SVM-Linear) was selected for further analysis because it yielded performance comparable to SVM with a radial kernel (SVM-Radial), while being substantially more computationally efficient. Seven of the nine methods (SVM, RF, GBM, RPART, PLS, KNN and NB) were trained and tuned using the caret package [19] v6.0-94 on R 4.3.3; ANN was tuned with the nnet package v7.3-19, and AdaBoost was implemented with scikit-learn [20] v1.9.1 (AdaBoostClassifier, SAMME algorithm), because the R ada package supports only binary classification. CellTypist v1.7.1 [3] was installed from PyPI on Python 3.11.14 and trained with celltypist.train. The Geneformer source code (commit 1f7fbae) was obtained from its Hugging Face repository (https://huggingface.co/ctheodoris/Geneformer) and imported directly from the cloned source on Python 3.12, with its dependencies installed from the repository’s requirements.txt (PyTorch v2.8.0 with CUDA 12.8, transformers v4.46.0); the Geneformer-V1-10M weights were downloaded from the same repository with huggingface_hub. In the present study, all methods were used to construct classification models on the GSE188515 dataset with their optimal hyperparameters, which were determined through exhaustive grid search with 10-fold cross-validation. Model performance was evaluated using accuracy and MCC. These measures were described in detail in our previous study [21] and calculated using in-house R scripts.

A unified data preprocessing workflow was implemented for all ten shallow-learning methods. First, library size normalization [8] was performed: all values in each column were divided by the total sum of that column, multiplied by 10,000, augmented with a pseudo-count of 1, and subsequently log2-transformed. Next, all 13 rows corresponding to mitochondrial genes were removed, followed by filtering out genes for which the proportion of cells with raw counts ≥ 2 was less than 1%. Thereafter, z-score standardization was applied: all values in each row were centered by subtracting the row-wise sample mean and then scaled by dividing by the row-wise sample standard deviation. Finally, PCA was performed, with the top 50 principal components retained as input for training. For the Geneformer-based models, raw counts from GSE188515 were converted to rank-value encoding using the official TranscriptomeTokenizer (model_version = “V1”): gene symbols were mapped to Ensembl IDs, the expression of each gene was normalized by its median across the pre-training corpus (gc30M dictionary), and the top 2,048 ranked genes per cell were used as input for training. Other training arguments were the official defaults (batch size 12, linear learning-rate schedule with 500 warm-up steps, weight decay 0.001, no frozen layers, random seed 42).

Following the default CellTypist procedure, the features were standardized (with_mean = True) and values greater than 10 were clipped to 10. A one-vs-rest L2-regularized logistic regression was fitted with the L-BFGS solver and a maximum of 1,000 iterations; stochastic gradient descent and mini-batch training were not used (use_SGD = False, mini_batch = False). Because scikit-learn ≥ 1.8 no longer supports the multi_class = “ovr” option used by CellTypist, the one-vs-rest classifier was reimplemented by fitting one binary logistic regression per class, which is equivalent to the original implementation. The regularization parameter C was tuned over 0.01, 0.1, 1, 10, 100 and 1000 according to the pooled out-of-fold accuracy, and C = 1 (the CellTypist default) was optimal. Predictions were made in “best match” mode, assigning each cell to the class with the highest probability; majority voting was not applied because it requires over-clustering of the expression data.

### Nankai GeneRank Classifier and SCPM-based clustering

Nankai GeneRank Classifier was constructed by fine-tuning the pre-trained Geneformer-V1 model and is accessible via a web-based interface at https://d2xdiv0mk15orl.cloudfront.net/. The v1 classifier was trained on 16 cell types comprising 10,784 cells, whereas the v2 classifier was trained on 29 cell types comprising 54,362 cells. SCPM-based clustering was implemented using Python 3.12.3; its workflow and comparison with Seurat clustering are detailed in **Supplementary File 1**. The normalized cc-scRNA783 dataset is available on the website, with its preprocessing described in **Supplementary File 1**. Both SCPM-based clustering and Seurat clustering processing for cc-scRNA783 are also detailed in **Supplementary File 1**.

To map bulk samples in the single-cell UMAP without altering the existing single-cell embedding, we employed the ingest function from Scanpy v1.12.4. Each bulk sample was mapped into this UMAP embedding as follows: its 15 nearest single cells were identified by Euclidean distance in clr space, and its initial position was calculated as the weighted average of their UMAP coordinates, with closer cells assigned larger weights. This position was subsequently fine-tuned through UMAP optimization. The cluster labels and UMAP coordinates of the single cells thus remained unchanged. As a control, the 783 single cells were projected using the same procedure to confirm that they returned to their original positions. Additionally, for each bulk sample, the cluster to which the majority of its 15 nearest single cells (by Euclidean distance in clr space) belonged was recorded as its nearest cluster, and its probabilities across the 29 pre-defined cell types were reported as its tissue-to-cell-type similarity profile.

## Conclusion

In the present study, we introduce an SCPM-based clustering method designed to streamline cell type identification using scRNA-seq data. The workflow comprises two steps: (i) SCPM generation by the Nankai GeneRank Classifier and (ii) a subsequent clustering step. Our new method enhances the resolution of cell type identification through clustering and ensures its accuracy through the classifier. As an additional key advantage, it enables the unified processing of bulk RNA-seq and scRNA-seq datasets for direct cross-modal comparison. Given that the present study focuses primarily on the feasibility of the proposed method, the diversity and number of cell populations in the training datasets are relatively limited, which may constrain the accuracy and resolution of cell type identification.

Leveraging SCPM-based clustering, we re-identified five cell populations in the cc-scRNA783 colorectal cancer dataset, namely ccEC, ccEpC, malignant ccEpC, cancer-like ccEpC, and CSCs. Building on this refined annotation, we further discovered a panel of candidate CSC marker genes. Re-analysis of the cc-scRNA783 dataset confirmed the previous finding [9] that no single previously published marker could effectively distinguish colorectal CSCs from other cell populations. Among the candidate markers identified here, PTPRO, IGFL4, MID1 [17], and LINC02563 [18] were more specifically expressed in CSCs than both the early reported markers (e.g., PROM1/CD133 and CD44) and the five recently reported markers [9] (NOTUM, SMOC2, BAMBI, PHLDA1, and TNFRSF19). Benefiting from the enhanced resolution of cell type identification achieved by our method, we further revealed that the sharp up-regulation of PTPRO occurs at the transition from cancer-like ccEpC to CSC—a finding that would be missed using classical methods.

## Authors’ contributions

Shan Gao conceived the project. Shan Gao and Xin Li supervised this study. Yanting Guo and Guangyou Duan performed programming. Guofeng Xu and Jia Chang downloaded, managed and processed the data. Chang Liu validated the results. Shan Gao drafted the main manuscript text. Shan Gao and Xin Li revised the manuscript.

## Data Availability Statement

All data used in the present study are provided in the Supplementary Files.

## Acknowledgments

We sincerely thank Professor Lin Liu and his former student Hua Wang from College of Life Sciences at Nankai University. This manuscript was submitted as a preprint on Sep. 15th, 2026 at https://www.biorxiv.org/content/10.64898/xxxxxxv1.

## References

[1] Gao, S., Data Analysis in Single-Cell Transcriptome Sequencing. Methods in Molecular Biology, 2018. 1754: p. 18.

[2] Satija, R., Farrell, J., Gennert, D. et al. Spatial reconstruction of single-cell gene expression data. Nat Biotechnol 33, 495–502 (2015).

[3] Domínguez Conde, C., Xu, C., et al. (2022). Cross-tissue immune cell analysis reveals tissue-specific features in humans. Science, 377(6607), eabl5197 .

[4] Theodoris, C. V., et al. (2023). Transfer learning enables predictions in network biology. Nature, 618(7965), 616–624.

[5] McInnes, L., Healy, J., Saul, N., & Grossberger, L. (2018). UMAP: Uniform Manifold Approximation and Projection. Journal of Open Source Software, 3(29), 861.

[6] Devlin, J., Chang, M.-W., Lee, K., & Toutanova, K. (2019). BERT: Pre-training of deep bidirectional transformers for language understanding. Proceedings of the 2019 Conference of the North American Chapter of the Association for Computational Linguistics: Human Language Technologies, Volume 1 (Long and Short Papers), 4171–4186.

[7] Vaswani, A., Shazeer, N., Parmar, N., Uszkoreit, J., Jones, L., Gomez, A. N., Kaiser, Ł., & Polosukhin, I. (2017). Attention is All you Need. Advances in Neural Information Processing Systems, 30.

[8] Zhenfeng Wu, Weixiang Liu, Xiufeng Jin, Haishuo Ji, Hua Wang, Gustavo Glusman, Max Robinson, Lin Liu, Jishou Ruan, Shan Gao. NormExpression: an R package to normalize gene expression data using evaluated methods. Frontiers in Genetics 2019,

[9] Hua Wang, Peng Gong, Tong Chen, Shan Gao, Zhenfeng Wu, Xiaodong Wang, Jie Li, Sadie L. Marjani, José Costa Sherman M. Weissman, Feng Qi, Xinghua Pan, Lin Liu. Colorectal Cancer Stem Cell States Uncovered by Simultaneous Single-Cell Analysis of Transcriptome and Telomeres. Advanced Science 2021, 8(8): 1–19.

[10] Fulong Ji, Yong Liu, Jinsong Shi, Chunxiang Liu, Siqi Fu, Heng Wang, Bingbing Ren, Dong Mi, Shan Gao and Daqing Sun. Single-Cell Transcriptome Analysis Reveals Mesenchymal Stem Cells in Cavernous Hemangioma. Frontiers in Cell and Developmental Biology 2022, 10: 1–8.

[11] Yaping Fang, Shan Gao, David Tai, C Russell Middaugh, Jianwen Fang. Identification of properties important to protein aggregation using feature selection. BMC Bioinformatics 2013, 14(1):314.

[12] Wanxing Wang, Jiawei Zhang, Guangyou Duan, Yiyao Zhang, Guofeng Xu, Zhi Cheng, Shunmei Chen, Xue Yao, Xin Li, Shan Gao. Coordinate Up-regulation of Both TK1 and TS Expression in Cycling Cells: Mechanistic Basis and Implications for Dual-Targeted Cancer Therapy. Biorxiv, 2026, 697331.

[13] Jiawei Zhang, Jingsong Shi, Liangge Wang, Xinjie Liu, Zemin Cao, Cihan Ruan, Guangzhi Ning, Shiqing Feng, Xue Yao, Shan Gao. Re-analysis of single-cell RNA-seq data reveals the origin and roles of cycling myeloid cells. Stem Cells 2024, 42(7): 593–606.

[14] Dai S, Wang Y, Liu H, et al. PTPRO represses colorectal cancer tumorigenesis and progression by reprogramming fatty acid metabolism[J]. Cancer Communications, 2022, 42(10): 1048–1067.

[15] Sun Y, Li Z, Zhang X, et al. Single-cell transcriptomic analysis deciphers heterogenous cancer stem-like cells in colorectal cancer and their organ-specific metastasis[J]. Gut, 2024, 73(3): 470–485

[16] Chen C Y, Yu H F, Han F Y, et al. Tumor-suppressive circRHOBTB3 is excreted out of cells via exosome to sustain colorectal cancer cell fitness[J]. Molecular Cancer, 2022, 21(1): 46.

[17] Zhao Z, Yang L, Ding Y Q, et al. Expression of MID1 in colorectal carcinoma and its prognostic significance[J]. World Chinese Journal of Digestology, 2012, 20(2): 113–118.

[18] Wang X C, Liu Y, Zhang M, et al. Identification of a lncRNA prognostic signature related to stem cell index and its significance in colorectal cancer[J]. Future Oncology, 2021, 17(23): 3087–3100.

[19] Kuhn M: Building Predictive Models in R Using the caret Package. J Stat Softw 2008, 28(5):1–26.

[20] Pedregosa et al. (2011). Scikit-learn: Machine learning in Python. JMLR, 12, 2825–2830.

[21] Shan Gao, Ning Zhang, Guangyou Duan Zhuoyang, Jishou Ruan, Tao Zhang. Prediction of function changes associated with single-point protein mutations using support vector machines (SVMs). Human Mutation 2009, 30(8):1161–1166.

